# Population genomics of yellow-eyed penguins uncovers subspecies divergence and candidate genes linked to respiratory distress syndrome

**DOI:** 10.1101/2025.10.20.683354

**Authors:** Joseph Guhlin, Janelle R Wierenga, Jordan Douglas, Puawai Swindells-Wallace, Hoani Langsbury, Trudi Webster, Melanie J. Young, Hendrik Schultz, Jordana Whyte, Bryony Alden, Thor T. Ruru, Leith Thomson, Jason van Zanten, Megan Abbott, Jim Watts, Harry S Taylor, Stuart Hunter, Yolanda van Heezik, Philip J. Seddon, Kerri Morgan, Lisa S. Argilla, Catherine E Grueber, Anna W Santure, Peter K Dearden, Jemma L Geoghegan

## Abstract

Yellow-eyed penguins (hoiho/takaraka, *Megadyptes antipodes*) are among the world’s rarest penguins and are regarded as a taonga (treasured) species in Aotearoa New Zealand. Since 2019, chicks on the New Zealand mainland have been affected by a deadly neonatal disease called respiratory distress syndrome (RDS), contributing to a decline to fewer than 143 breeding pairs. To investigate the putative genetic basis of this disease, we generated high-quality whole-genome data from 249 individuals spanning the species’ range, including from the New Zealand mainland (Northern range) and subantarctic Enderby and Campbell Islands (Southern). Population genomic analyses unexpectedly revealed three deeply divergent lineages with negligible gene flow, consistent with recognition of three distinct subspecies.

Phylogenetic divergence dating suggests that these splits predate human arrival by several millennia, with the Northern lineage diverging from the Southern populations 5-16 ka. Genome scans for local adaptation revealed regions of strong differentiation, and genome-wide association analyses identified candidate immune and respiratory genes linked to RDS. In partnership with Ngāi Tahu, who hold indigenous guardianship over yellow-eyed penguins, we recommend recognition of three subspecies, urgent conservation action for the critically small and rapidly declining Northern subspecies, and the need for immediate population size and trend assessments for Auckland and Campbell Island populations.

## Introduction

Yellow-eyed penguins (*Megadyptes antipodes*) are an endangered species endemic to Aotearoa New Zealand^1^. They are regarded as taonga (treasured) by Māori, the indigenous peoples of New Zealand, who named them *hoiho* or *takaraka*. As one of the rarest penguin species in the world, yellow-eyed penguins serve as an important indicator of ecosystem health, with their population trajectories reflecting broader ecosystem change^1-3^. Protecting yellow-eyed penguins is therefore critical for biodiversity conservation and maintaining the integrity of coastal ecosystems. They are also an icon of the regional wildlife tourism industry, significantly contributing to the local economy^4^. Their decline therefore represents a biodiversity crisis as well as a cultural and economic loss.

Yellow-eyed penguins have been delineated into two management groups: the *Northern* population found on the New Zealand mainland (throughout Te Waipounamu/South Island and Rakiura/Stewart Island), and the Southern population distributed across Maukahuka/Motu Maha/Auckland Islands, including Enderby Island, as well as Motu Ihupuku/Campbell Island (Figure 1a). Today, fewer than 3,000 individuals remain, with c.143 breeding pairs remaining within the Northern distribution (Department of Conservation, unpublished data), and fewer than 20% of Northern chicks surviving to adulthood^5^. As the sole-surviving species of the *Megadyptes* genus, the yellow-eyed penguin is not only Endangered^6,7^, but at risk of extinction within its northern range within the next two decades due to threats of habitat loss, dietary change^8-11^, disease^12-18^, bycatch^19-21^, and climate change^22,23^.

**Figure 1.**
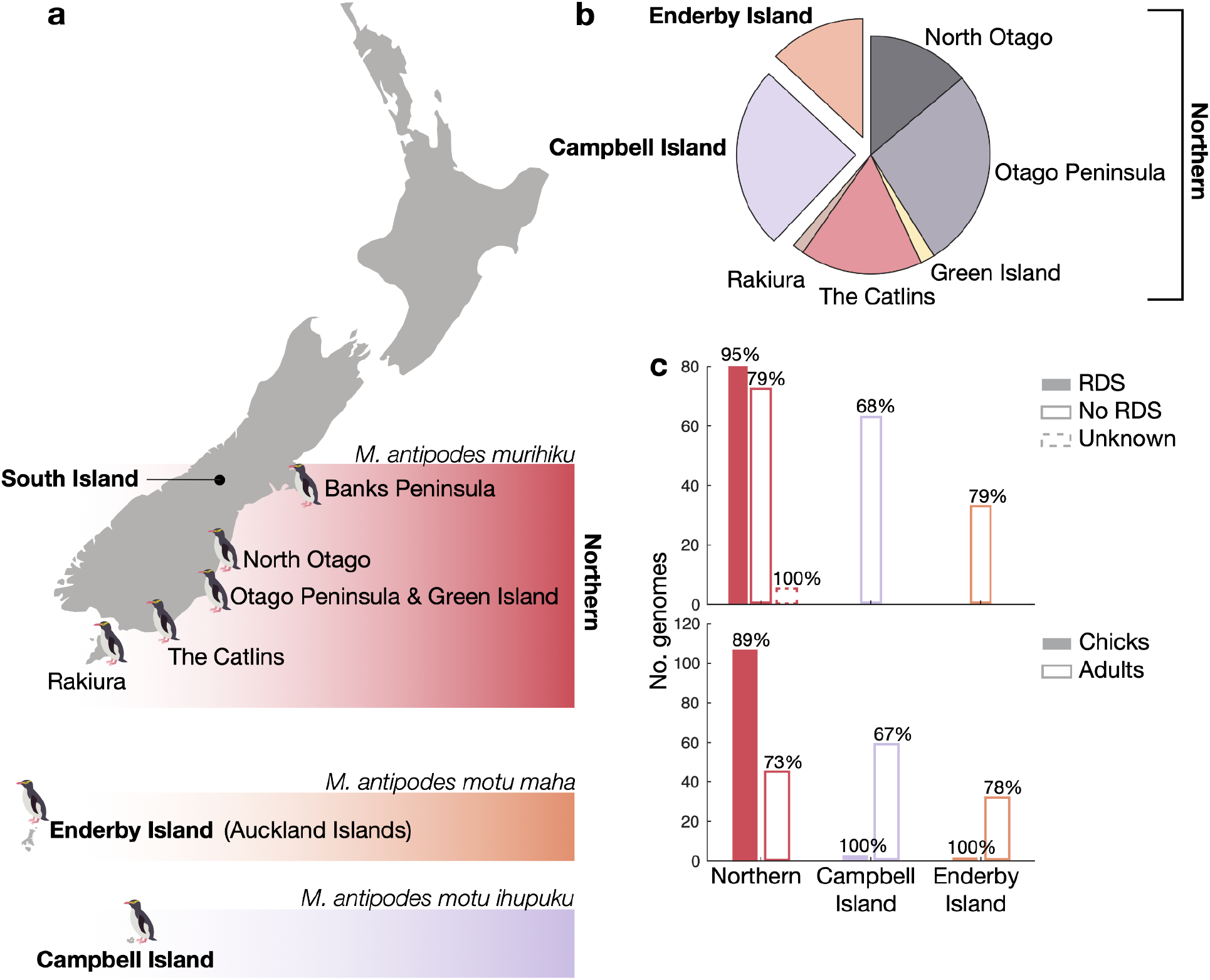
(**a**) Map of Aotearoa New Zealand indicating the geographic distribution of yellow-eyed penguins and proposed new subspecies designations. (**b**) Pie chart showing the proportion of individuals sampled from each location. Note that no samples were obtained from individuals at Te Pātaka-o-Rākaihautū/Banks Peninsula and that only Enderby Island individuals were sampled from Maukahuka/Auckland Islands. (**c**) Number of samples from each population broken down by respiratory distress syndrome (RDS) diagnosis and age class, noting the percentage of samples in which *Yellow-eyed penguin gyrovirus* (YPGV) were detected via PCR above each bar.

Since 2019, a deadly neonatal respiratory disease known as respiratory distress syndrome (RDS) has presented an additional threat to the persistence of the Northern population. RDS results in severe breathing difficulties, with necropsies revealing lung congestion and haemorrhage along with lymphoid depletion^24^. The causative agent is likely a non-enveloped DNA virus in the *Anelloviridae* family known as *Yellow-eyed penguin gyrovirus*, or YPGV^24^. In contrast, despite the endemicity of YPGV in all regions, the Southern population shows no apparent clinical signs consistent with RDS. Whether the Northern population is genetically more susceptible to developing RDS, or the differences in disease presentation are due to other causes (such as environmental variables), remains an open question requiring species-wide genomic studies.

Previous genetic studies have suggested that yellow-eyed penguins migrated to their Northern range c. 500 years ago, after the extinction of the closely related *Megadyptes antipodes waitaha*, coincident with human arrival on the mainland^25-28^. In this recent-immigrant model, the mainland population is viewed as having established via a peripheral northward range expansion of the species, and is potentially maladapted to warmer conditions. Under this hypothesis, conservation strategies for the Northern population could involve replacement, translocation, or genetic rescue from Southern populations, noting that at the present time, no such strategies have been proposed^29^. Yet, before such interventions can be realistically considered, the genetic basis of the populations’ apparent differential susceptibility to developing RDS must be resolved, as this remains a key knowledge gap that could directly inform conservation decision-making.

In this study, deep sequencing of 249 yellow-eyed penguin genomes further resolved the species’ demographic history and allowed us to investigate the genetic basis of susceptibility to developing RDS. We combined phylogenetic molecular dating, genome-wide association studies^30^, selective sweep scans^31^, and ancestral recombination graph (ARG) inference^32,33^ to capture both long-term and recent evolutionary dynamics. Among these, ARGs were particularly suited to our aims, as they enabled the estimation of the time to the most recent common ancestor both within and between populations. This fine-scale approach offers substantially greater resolution of divergence, migration, and adaptation over the last few dozen generations than traditional population genetic methods^34,35^.

Through this framework, we have revealed the presence of three genetically distinct populations of yellow-eyed penguins. These lineages are characterised by deep divergence, minimal genomic evidence of ongoing migration, and signatures of local adaptation. The scale of genomic differentiation strongly supports their recognition as three genomically and geographically distinct subspecies, with the split between the Northern and Southern populations occurring much earlier than previously considered^25^. We have identified genetic variants that underlie differential responses to infection with YPGV, highlighting the role of host genetics in disease outcomes. Given the cultural and economic significance of yellow-eyed penguins, our finding that the Northern, Enderby Island, and Campbell Island populations represent three distinct subspecies underscores the urgent need for recognition and separate management strategies in the face of population declines. Without immediate conservation action, each lineage faces risk of localised extinction.

## Results

### Generating yellow-eyed penguin genomes and population sequencing

We assembled reference genomes for two individuals from each management unit, one Northern and one Southern, and generated whole-genome resequencing data for 253 yellow-eyed penguins (249 of which were successful) to investigate genetic associations with respiratory distress syndrome (RDS), characterise population structure, and estimate divergence times among populations. Sampling, by venepuncture, comprised individuals from the Northern population from post-mortem samples, and from Enderby Island in the Auckland Islands, and Campbell Island (Figure 1). Approximately 60% of genomes were derived from the Northern population; of these, approximately half (51%) had a confirmed diagnosis of RDS based on gross postmortem examination, providing a robust case-control dataset for association testing. PCR revealed that *Yellow-eyed penguin gyrovirus* (YPGV), while not present in all samples, was present across all sampled regions and age classes (Figure 1). The ubiquity of YPGV detection highlights its widespread circulation within the species and provides important context for evaluating potential interactions between host genetics, viral infection, and disease outcomes.

The Northern reference genome (A9) assembled to 1.358 Gbp across 595 contigs, with an N50 of 22.868 Mbp, while the Southern reference genome (C90) was 1.278 Gbp across 30,923 contigs, with an N50 of 78,675 kbp. Repeat masking resulted in 16.51% masked sequence of the A9 genome and 13.72% of the C90 genome. Genomic completeness, assessed using BUSCO^36^, indicated that A9 was highly complete at 99.5%, with only 0.4% missing and 0.6% duplicated. Our C90 genome assembly was just 78.6% complete, with 6.7% missing and 0.8% duplicated. For A9 and C90, we identified 63,026 and 71,337 protein-coding genes, respectively. Protein-level BUSCO analysis resulted in 97.7% complete, with 1.8% missing for A9 gene predictions, while C90 was 78.1% complete, with 7.7% missing. Full details are available in Table S1. Population samples were sequenced to an average sequencing depth of 28.12x (range 3.01x - 67.58x, standard deviation = 5.82x) coverage (Table S2).

### Variant Statistics

After removing three individuals due to low sequencing output and one individual due to unexpectedly high heterozygosity suggestive of a sample mixture, we mapped reads to the A9 genome and called an all-samples single nucleotide polymorphism (SNP) set of 2,251,578 biallelic SNPs across 249 individuals. Filtering for minor allele count (MAC) of ≥2 resulted in 1,855,733 SNPs. Per population counts, with individual population-level filtering, were 1,444,663 for the Northern population, 1,058,683 for Enderby Island, and 1,232,200 for Campbell Island. The Northern population had lower median heterozygosity and higher median inbreeding than the other two populations, while a paucity of rare alleles in the folded allele frequency spectrum suggest that yellow-eyed penguins from Enderby Island may have had a recent bottleneck event (Figure S1-S3). We downloaded ancient DNA sample reads generated from two closely related extinct penguins; *Megadyptes antipodes richdalei* and *M. a. waitaha*^37^, and aligned these to our A9 reference genome. We identified 349,778 SNPs for *M. a. richdalei*, and 28,435 SNPs for *M. a. waitaha*.

### Ancestral recombination graphs

To reconstruct the complete ancestral history of the 249 sampled individuals, we inferred their Ancestral Recombination Graphs (ARGs), which model both coalescence and recombination events along the genome. The resulting ARGs provide a full representation of the historical genealogical relationships among all individuals in our three subspecies, offering greater insight beyond the scope of a single consensus tree, thus capturing complex patterns of genomic inheritance. In total, we generated 686,020 local trees, each describing the ancestry of a specific genomic segment. Together, these trees form a genome-wide genealogical record that underpins downstream analyses, providing the framework for estimating coalescence times, detecting selective sweeps, and accounting for genetic relatedness in the genome wide association study (GWAS)^38^.

### Population differentiation, structure, and migration

To investigate the population genomic structure of yellow-eyed penguins, we analysed genome-wide variation across individuals from the Northern and Southern range. The Northern population formed a distinct genetic cluster (Figure 2), with no evidence of contemporary migration or admixture with Southern populations, with some levels of migration between the Northern locations (Figure S4, S5). Importantly, individuals with and without RDS did not cluster separately, indicating that RDS status is not confounded by population structure.

**Figure 2.**
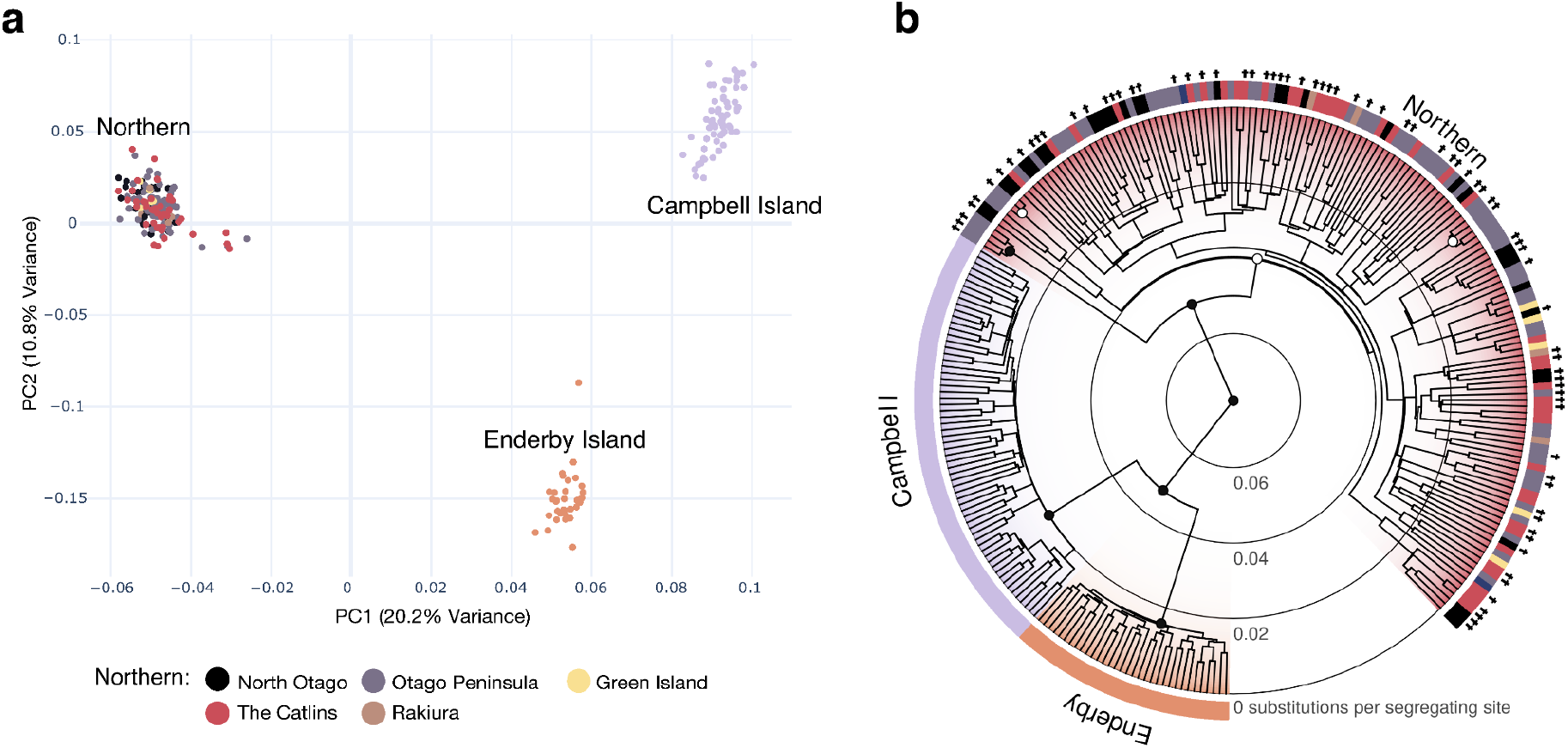
The three populations are clearly delineated by their genomics, but there is no evidence of any genetic structure within each of the populations. (**a**) The first two principal components were calculated from 2,251,578 nuclear SNPs. Sampling locations follow the same colour scheme as Figure 1. (**b**) Evolutionary relationships estimated by the SNAPPER^39^ multispecies coalescent model from 200 randomly sampled SNPs. Black circles indicate clades with over 95% posterior support, and white denotes clades with 50 to 95% support. Individuals with an RDS diagnosis upon gross post-mortem are indicated with a cross, while the remaining did not have RDS.

The Southern population was subdivided into two genetically discrete groups corresponding to Campbell Island and Enderby Island. Estimates of gene flow between these populations indicated negligible levels of connectivity, consistent with long-term isolation and independent demographic histories. These findings support recognition of three evolutionarily distinct lineages of yellow-eyed penguins (Figure 2).

Following consultation with Ngāi Tahu, the principal iwi (tribe) of the South Island, New Zealand, who are kaitiaki (guardians) for the species, we propose the designation of three subspecies of yellow-eyed penguins that reflects their evolutionary divergence. Specifically, we recommend recognition of: (i) *Megadyptes antipodes murihiku* (hoiho murihiku) for the Northern population (Murihiku referring to the southern region of the South Island); (ii) *Megadyptes antipodes motu maha* (hoiho motu maha) for Enderby Island within the Auckland Islands (Motu Maha being one of the Māori names for the archipelago); and (iii) *Megadyptes antipodes motu ihupuku* (hoiho motu ihupuku) for Campbell Island (Motu Ihupuku being the Māori name for the island).

### Yellow-eyed penguin divergence

To understand the timing of divergence events we estimated a dated phylogeny of the *Megadyptes* and their close relatives (Figure 3). These relatives were the extinct *Megadyptes antipodes waitaha* and *M. a. richdalei*, plus representatives from the *Eudyptes, Eudyptula*, and *Spheniscus* genera. Genomic sequences of *M. a. waitaha* and *M. a. richdalei*, and estimated divergence times across penguin species (Table 1), were generated by Cole et al.^37^, with additional species from Pan et al.^40^ and Li et al.^41^. We estimated the species phylogeny from 50 genomic loci across a total of 264 penguins (249 of which from our population resequencing). This was performed under a multispecies coalescent model in StarBeast3^42^, using both a strict and relaxed clock model. The strict clock assumes a constant rate of evolutionary change throughout time, while the relaxed clock allows the clock rate to vary independently across lineages, and is often thought to be more realistic over longer timescales. Moreover, by taking a multilocus approach (StarBeast3^42^ here, and SNAPPER^39^ in the analysis shown in Figure 2), we avoid statistical biases that arise from concatenating loci^43,44^.

**Table 1.**
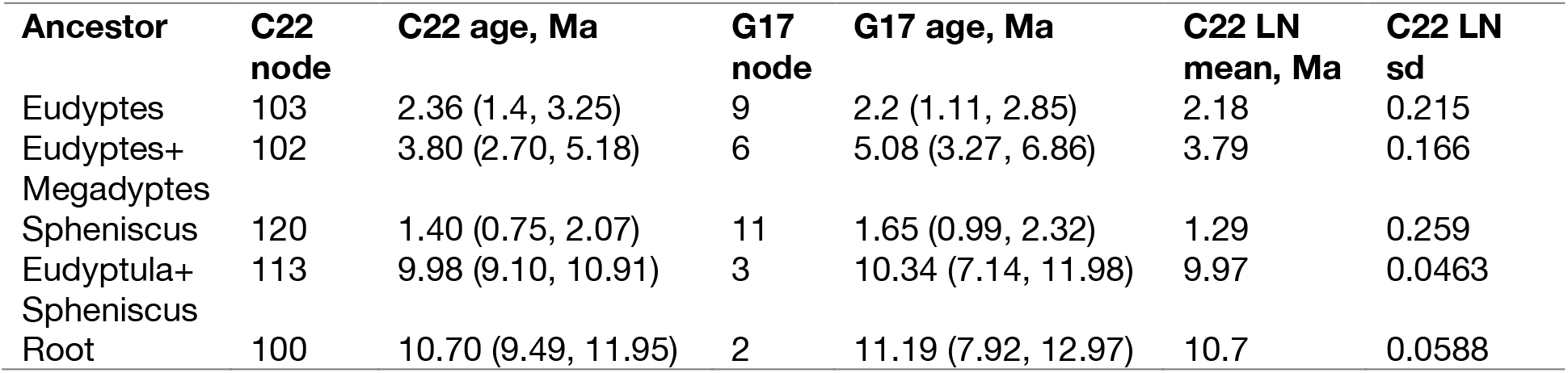
Secondary calibrations used in our StarBeast3 analysis. For each calibration point (shown as diamonds on Figure 3), we report the ages estimated by Cole et al.^37^ (C22) and Gavryushkina et al.^45^ (G17). These ages are expressed as means (and 95% credible intervals), with the node index within the original studies indicated. Our log-normal prior distributions each have a 95% interquartile interval equal to the 95% credible intervals of C22. These log-normal distributions are expressed with their mean in real space (LN mean) and standard deviation in log space (LN sd).

| Ancestor | C22 node | C22 age, Ma | G17 node | G17 age, Ma | C22 LN mean, Ma | C22 LN sd |
| --- | --- | --- | --- | --- | --- | --- |
| Eudyptes | 103 | 2.36 (1.4, 3.25) | 9 | 2.2 (1.11, 2.85) | 2.18 | 0.215 |
| Eudyptes+ Megadyptes | 102 | 3.80 (2.70, 5.18) | 6 | 5.08 (3.27, 6.86) | 3.79 | 0.166 |
| Spheniscus | 120 | 1.40 (0.75, 2.07) | 11 | 1.65 (0.99, 2.32) | 1.29 | 0.259 |
| Eudyptula+ Spheniscus | 113 | 9.98 (9.10, 10.91) | 3 | 10.34 (7.14, 11.98) | 9.97 | 0.0463 |
| Root | 100 | 10.70 (9.49, 11.95) | 2 | 11.19 (7.92, 12.97) | 10.7 | 0.0588 |

**Figure 3.**
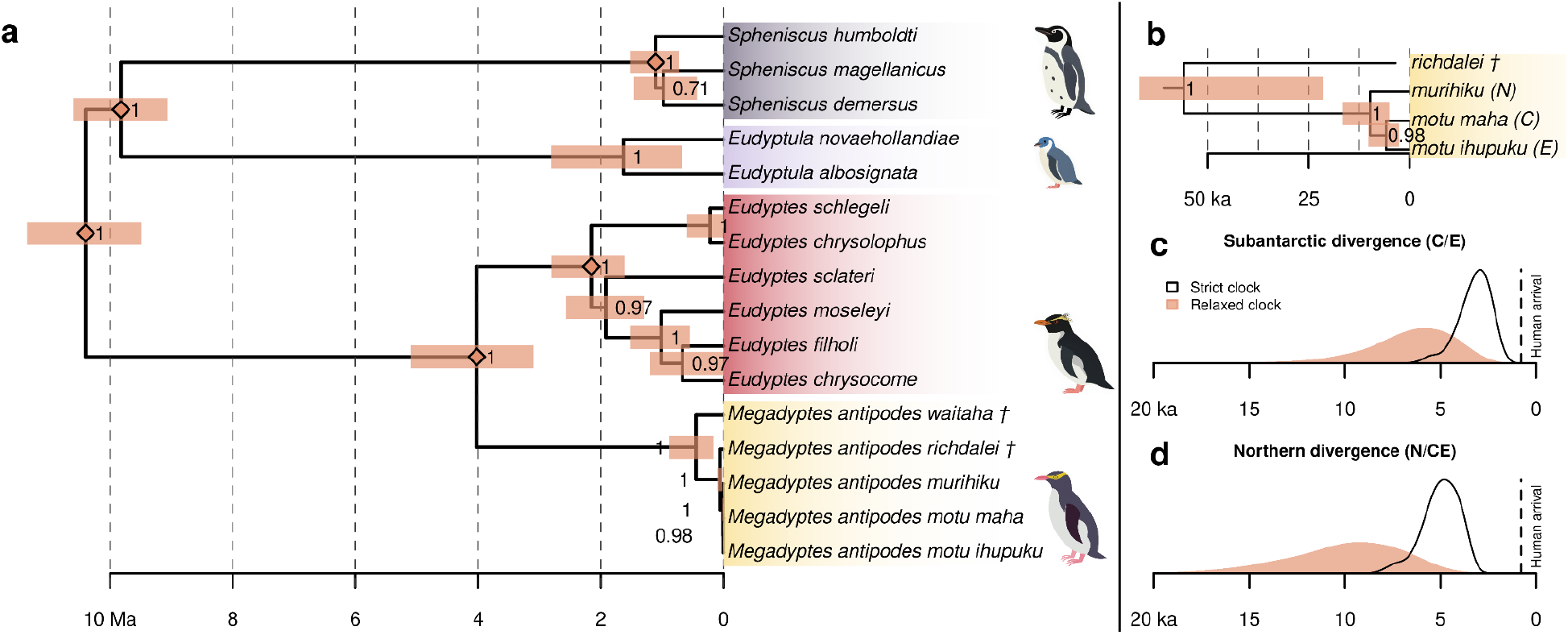
Molecular dating analysis under a multispecies coalescent model (StarBeast3^42^) with a relaxed clock. (**a**) Internal nodes show the posterior clade support and 95% highest density posterior interval of divergence time. The five nodes selected for secondary calibration are indicated with diamonds^37^. (**b**) A close-up of the *Megadyptes antipodes* clade in (a). The posterior distributions of the two recent *Megadyptes antipodes* divergence events are shown for (**c**) Campbell Island (C) from Enderby Island (E), and (**d**) Northern population (N) from Campbell and Enderby Islands.

Results suggest that the extant *Megadyptes* subspecies split much earlier than previously indicated. While previous data placed the split around the 16th century^25^, both of our models suggest it occurred long before the first peoples arrived in New Zealand (ca. 1250 CE). We found that the Northern-Southern divergence happened first, and the Campbell-Enderby Islands divergence afterwards. The strict clock places the first split 5,000 years ago, with a 95% credible interval of (3,300 – 6,800), whereas the relaxed clock pushes this back even further to 10,000 years (4,900 – 16,000). The Southern (subantarctic) subspecies were estimated to have split 3,200 years ago (1,700 – 4,800) under a strict clock and 6,700 years ago (2,600 – 11,000) under a relaxed clock. Had we used the secondary calibrations from Gavryushkina et al.^45^, rather than Cole et al.^37^, these ages would likely have been slightly older (Table 1). The molecular clock rates were estimated as 1.75 × 10^-9^ substitutions per site per year (1.42 × 10^9^ – 2.08 × 10^9^) under the strict clock, and 1.55 × 10^-9^ (1.06 × 10^9^ – 2.00 × 10^9^) for the relaxed clock. Evolutionary change was slightly faster under the strict clock, which explains why its divergence dates were more recent. These are both slower than the yellow-eyed penguin clock rates by Cole et al.^37^ (at 7 × 10^-9^), which may reflect different criteria for locus inclusion or different evolutionary models.

### Population trajectories

To find regions undergoing differential selection between subspecies, we used two approaches: (i) Cross-population extended haplotype homozygosity test (XP-EHH), and (ii) time to the most recent common ancestor (TMRCA) ratio. XP-EHH is pairwise, comparing between subspecies, with high absolute values indicating selective sweeps, and the sign of the statistic indicating which subspecies the sweep occurred in^46^. Between penguins sampled from Campbell and Enderby Islands, we detected 159 significant regions (|XP-EHH| >= 5.0), between those sampled from Campbell Island and the Northern population we found 243 regions; and between those from the Northern population and Enderby Island we found 145 regions (Figure 4).

**Figure 4.**
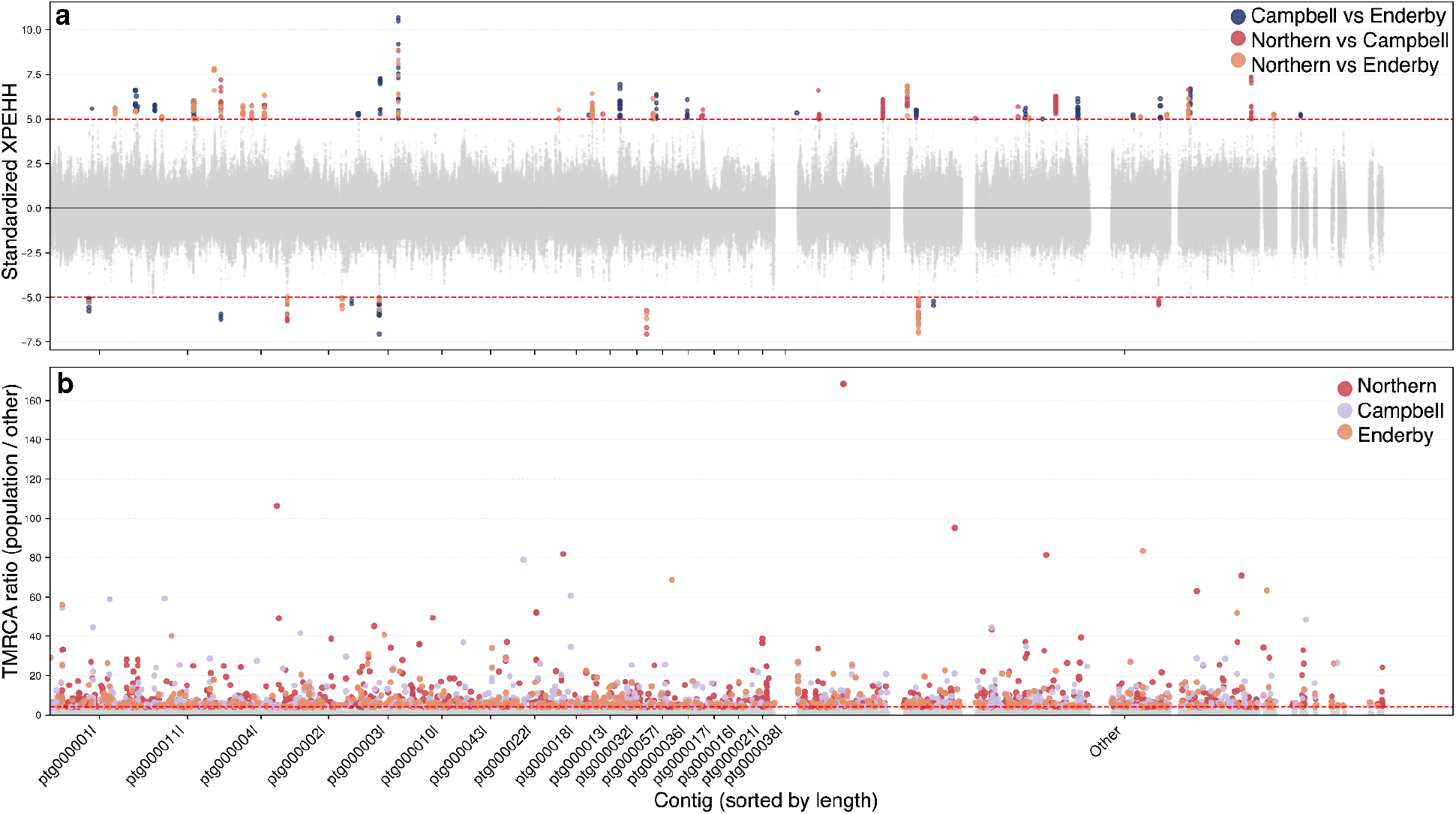
Genomic regions of divergence between Northern population, Campbell Island, and Enderby Island subspecies. (**a)** Between-population extended haplotype homozygosity (XP-EHH) analysis per subspecies pair; high absolute values indicate a selective sweep in the first subspecies relative to the second, low absolute values represent the opposite. (**b**) Time to most recent common ancestor (TMRCA) ratio, high values represent regions per subspecies where the TMRCA is young compared to the other subspecies, also indicative of a recent selective sweep in that subspecies.

The TMRCA ratio analysis identifies regions of the genome within each subspecies that have much younger TMRCA, indicative of subspecies-specific selective sweeps^31^. By looking only at regions in the top 0.1% quantile, we identified nine regions in yellow-eyed penguin genomes from Campbell Island making up 76.955 Mbp, three in those from Enderby Island (133.541Mbp), and 11 in the Northern range (137.801Mbp). The Northern subspecies showed significant gene enrichment for gene ontology (GO) categories for response to temperature stimulus, cold, nutrient levels, and salt stress (Bonferroni-corrected *p* < 0.05; Table S3). Yellow-eyed penguins from Campbell Island showed only two significant GO enrichment terms, dopamine transport and catecholamine transport (Table S4), while none were significant for those sampled on Enderby Island (Table S5).

Given the unique disease structure, where the virus is present in all three subspecies, yet clinical disease apparently only manifests in individuals from the Northern subspecies, we determined whether recent selective sweeps might differentiate between the Southern subspecies (i.e., those from Enderby and Campbell Islands) and the Northern, highly vulnerable subspecies. To do so, we compared TMRCA ratios for Southern versus Northern, and found younger TMRCAs in the Southern populations for cilium-related GO terms, including cilium beat frequency and movement, following Bonferroni correction. Cilia are microscopic, hair-like organelles that line the respiratory tract and provide a physical defense against inhaled pathogens. Given that nearby SNPs in the genome are non-independent, a Bonferroni correction is likely to be overly stringent, and it is warranted to examine other statistically significant signals excluding Bonferroni correction for additional biological insight. Notably, the eighth-ranked term was ‘myeloid dendritic cell differentiation’, which links the innate and adaptive immune systems, suggesting potential divergence in immune cell maturation pathways^47^. Several additional cilia-related processes were also elevated, further supporting a role for epithelial defense mechanisms in mediating resistance to respiratory infection (see Table S6 and S7 for the complete table of GO terms).

### Genome-wide association disease study

We identified variants associated with RDS status and YPGV presence (Figure 5, Tables S8 and S9). Small population GWAS studies are often ‘noisy’ but result in biologically meaningful peaks^48-50^ and experimentally validated phenotypic change^51-53^. Points of interest and association in GWAS are determined primarily by ‘peaks’ on a Manhattan plot (Figure 5). The associated marker is not expected to be the causative mutation, but a nearby mutation. Thus, we searched for genes within 20 kbp flanking top GWAS hits, identifying potential candidates for further study using both statistics and a literature search. Our analyses are well-calibrated for RDS status with λGC of 1.075, but we note inflated statistics for GV Status with λGC of 1.229 (QQ Plots, Figure S6).

**Figure 5.**
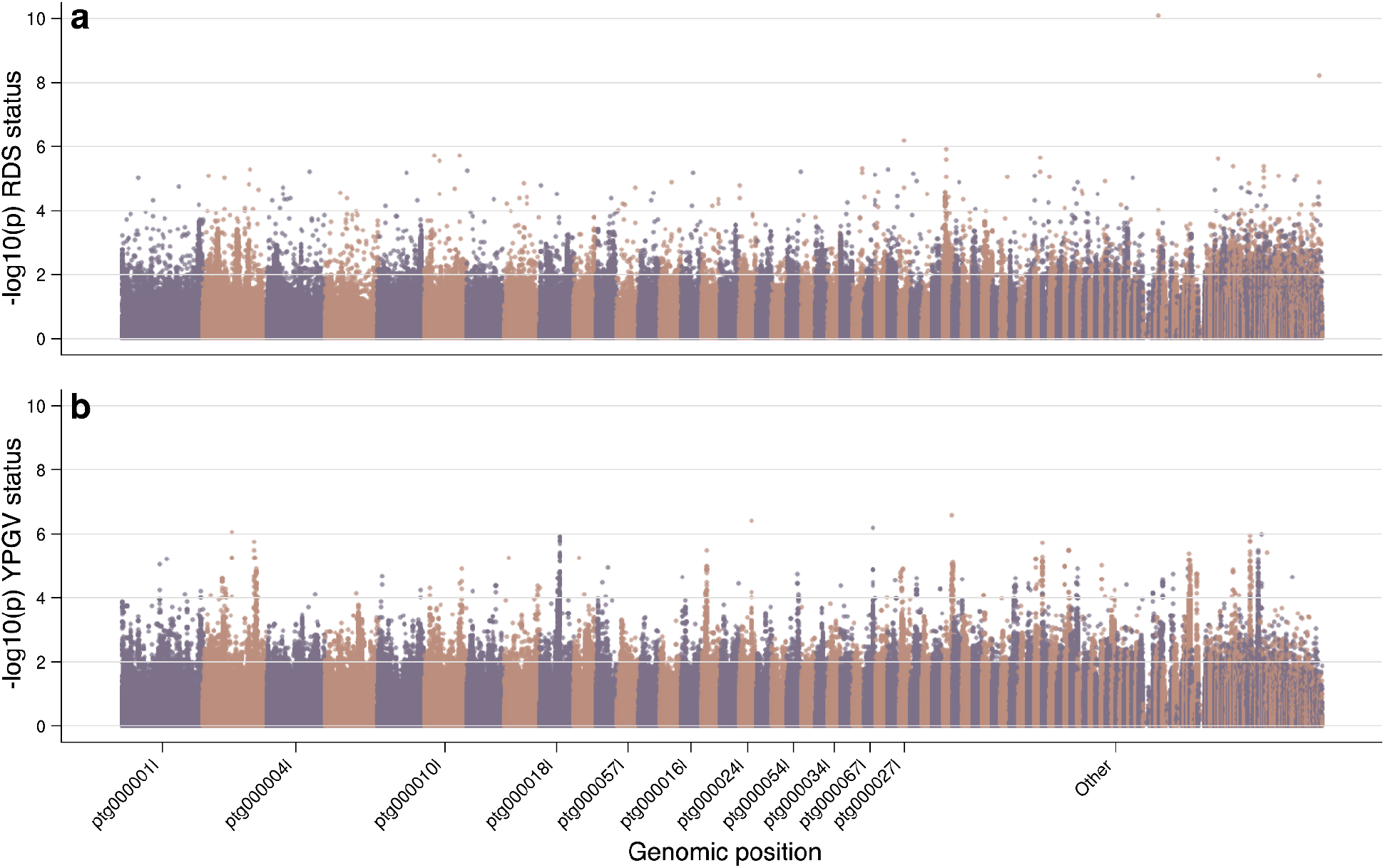
Genome Wide Association Study results for (**a**) respiratory distress syndrome (RDS) and (**b**) *Yellow-eyed penguin gyrovirus* (YPGV) presence phenotype, with *p*-values transformed with -log10p. Contigs are ordered in descending size. Colours alternate for each contig.

Our top GWAS signal, although lacking a pronounced peak likely due to the small contig size, corresponds to Pericentrin for RDS status. Pericentrin encodes a key centrosomal scaffolding protein essential for microtubule nucleation and organisation of the pericentriolar material (PCM). Viruses frequently co-opt this host machinery to facilitate dynein-dependent trafficking of viral particles along microtubules toward the nucleus, as well as to remodel the PCM during replication^54,55^. Pericentrin involvement here suggests that cytoskeletal and vesicular transport pathways may play a role in the viral entry or replication processes underlying RDS pathology. Another notable candidate is Anoctamin-6 (ANO6), a calcium-activated phospholipid scramblase. Scramblases regulate membrane asymmetry and have been shown to be exploited by diverse viruses, including coronaviruses and filoviruses, to mediate membrane fusion and budding events^56,57^. The potential association of ANO6 with RDS implicates altered host membrane dynamics or viral egress mechanisms in disease susceptibility.

Finally, gyrovirus presence showed a strong statistical association with High Mobility Group Box 1 (HMGB1), a DNA-binding protein that acts as an alarmin during infection. HMGB1 movement from the nucleus to the cytoplasm marks the onset of antiviral inflammation, activating immune signaling pathways that drive the production of pro-inflammatory cytokines. In poultry, HMGB1 is upregulated during severe viral infections such as *Newcastle disease virus*, where it amplifies the host inflammatory response^58^. This link may reflect an excessive or maladaptive immune response to gyrovirus infection in affected yellow-eyed penguin chicks.

Together, the convergence of these associations underscores the likely multifactorial and immune-modulated nature of RDS in yellow-eyed penguins, rather than a single gene or viral cause.

## Discussion

### Divergence timing of yellow-eyed penguins

Our analyses indicate that Northern and Southern yellow-eyed penguins comprise three distinct, isolated subspecies with no genetic evidence of migration or gene flow. Using a relaxed molecular clock with secondary calibrations from Cole et al.^37^, we estimate that the Northern subspecies diverged from the Southern group between 5,000 – 16,000 years ago, while the Campbell Island and Enderby Island subspecies diverged more recently, between 3,000 – 11,000 years ago (Figure 3). These divergence events likely pre-date human settlement in New Zealand (∼1250 CE). One possible interpretation is that yellow-eyed penguins first expanded northward from the subantarctic several thousand years ago, followed by a subsequent separation between individuals from Campbell and Enderby Islands. While the ancestral range of extant *Megadyptes* remains uncertain, the divergence times of these subspecies temporally align with a sequence of post-glacial environmental events^59^.

Earlier work by Boessenkool et al.^25^, based on ancient DNA from bones and DNA from modern samples, proposed a much more recent arrival of yellow-eyed penguins on the South Island following human-mediated extinction of *Megadyptes antipodes waitaha*. However, their initial inference relied on a single mitochondrial locus and an absence of *Megadyptes antipodes murihiku* bones in the limited pre-settlement material examined, much of which came from archaeological rather than natural contexts (the latter sites being beyond the endemic range of the extant *murihiku*^60^). Later analyses by the same group^26^ provided evidence for long-standing separation of Northern and Southern populations, aligning more closely with our genome-wide findings.

Taken together, the available genomic data are consistent with the view that *Megadyptes antipodes murihiku* have been present on the South Island of New Zealand for several thousand years. Additional fossil discoveries predating 1000 CE would help further resolve this history.

### Designation of three subspecies and conservation implications

Our genomic analyses support the recognition of three subspecies of yellow-eyed penguins: *Megadyptes antipodes murihiku* (Northern), *M. a. motu maha* (Enderby Island) and *M. a. motu ihupuku* (Campbell Island). The estimated divergence times, together with an absence of contemporary gene flow, indicate that these subspecies have been isolated for several thousand years and should be regarded as independent evolutionary lineages. Evidence of independent signals of selection in each subspecies, with regions enriched for genes relating to nutrition, stress response, temperature, and salt adaptation, adds further support to this conclusion.

This finding has direct conservation implications. The critically small population of the Northern subspecies, comprising approximately 143 breeding pairs and undergoing rapid decline due to dietary change, malnutrition, predation, bycatch, and diseases impacting survival and productivity including respiratory distress syndrome (RDS), cannot be supplemented through genetic rescue from other subspecies without the risk of outbreeding depression^61^. Instead, *M. a. murihiku* warrants urgent conservation attention due to the exacerbated rate of population decline^1^. Likewise, increased research and monitoring to better understand population size, trends, and threat mitigation options are required for Auckland Islands and Campbell Island subspecies, especially given the evidence for population instability^62^. Recognition of three distinct lineages underscores the need for tailored conservation strategies across the species’ range^63^.

### Genetic associations with respiratory distress syndrome

To investigate potential host genetic factors underlying susceptibility to RDS, we ran a selection scan analysis between the Northern and Southern subspecies, and conducted a genome-wide association study (GWAS) within the Northern subspecies. Our analysis indicated signals of selection (via a very recent time to most recent common ancestor) in the Southern subspecies at genomic regions enriched for genes related to cillial function and immunity. In the Northern subspecies, the GWAS revealed significant associations between RDS status and loci linked to genes with known roles in immune function and respiratory health in other avian taxa. These findings suggest there may be a genetic basis to the differential RDS susceptibility between Northern and Southern subspecies, but this remains to be experimentally verified. Further, higher inbreeding levels and lower heterozygosity in the Northern subspecies indicate genomic signatures of the ongoing population contraction. Long-term survival of this subspecies is therefore reliant on ongoing work to manage regional threats to population viability. This will include improved characterisation of the progression from viral infection to RDS and to develop effective control strategies for the virus to reduce chick mortality.

## Methods

### Ethics

Tissue samples were collected from routine post-mortems conducted under a Department of Conservation Contract in accordance with Animal Ethics Approval MUAEC Protocol 21/42 from Massey University and the New Zealand Department of Conservation Permit Authorisation Number 94843-FAU and 94902-FAU. Blood samples were collected by the Department of Conservation for species management purposes, mandated by Strategic Priority 2 of Te Mahere Rima Tau^29^ and authorised by the Department of Conservation Animal Ethics Committee AEC 405 and AEC 467. This project was undertaken in collaboration with the Department of Conservation and Te Rūnanga o Ngāi Tahu, the main iwi (tribe) of the South Island of New Zealand and who hold kaitiaki (guardianship) over yellow-eyed penguins and their data.

### Sample collection and DNA sequencing

Between 2021 and 2024, 253 yellow-eyed penguin samples were collected, comprising 157 (62%) from the Northern population and 96 (38%) from the Southern population. Samples were collected from North Otago, Otago Peninsula, Green Island, the Catlins, Rakiura, and Whenua Hou (Northern) as well as Enderby Island in the Auckland Islands and Campbell Island (Southern). Of the samples from the Northern population, 80 (51%) had RDS determined by gross post-mortem, and 76 (95%) of this diseased group tested positive, upon PCR, for YPGV (for PCR methods see Wierenga et al.^24^). Among the remaining healthy yellow-eyed penguins from the Northern population, 57 (79%) were YPGV positive. While RDS was absent from the Southern population, 69 (72%) had a YPGV infection.

Samples included either post-mortem liver tissue or blood. Liver tissue was macerated into small pieces by scalpel blade, added to Buffer ATL, and left to dissolve in a heat-block.

Approximately 1ml blood was collected by venepuncture from the metatarsal vein in lithium heparin green-topped tubes, plasma separated in red-topped tubes with no additive, or on filter paper blood cards. DNA was extracted using the Qiagen DNeasy Tissue and Blood kit (Qiagen) following the manufacturer’s instructions and DNA was subject to the Illumina DNA Prep M for library preparation and DNA sequencing on the NovaSeq X platform, targeting 20x sequencing read depth.

DNA from these extractions were also used for the production and assembly of two long read genome assemblies, one from a Southern population member (C90) and one from a Northern sample (A9). DNA libraries for Oxford Nanopore sequencing were produced using a Sequencing by Ligation Kit (version 14). Each sample was loaded on a P2 Solo Flow cell and data collected on a P2 Solo Nanopore device. For each sample, the library was recovered, and the flowcell washed multiple times to produce sufficient data for genome assembly.

Basecalling from this data was carried out using Dorado v. 0.9.0 (Oxford Nanopore Technologies).

### Assemblies and annotation

Our two assemblies were built using hifiasm v.0.25.0-r726^64^. Repeat finding was performed using REPRise v.1.0.1^65^ on the A9 assembly, and masking performed with RepeatMasker v4.2.1^66^ using the library generated from REPRise v1.0.1. BUSCO v6.0.0^36^ was used to report on genome completeness^36^. Gene annotation was performed using GALBA v1.0.11^67^ with Uniprot Swiss-Prot as the seed protein sequences, then BUSCO was repeated in protein/annotation mode^68^. Eggnog-Mapper v.2.1.12^69^ was used to associate genes with putative function, including Gene Ontology terms.

### ProgressiveCactus

Reference genomes existing for any Aequornithes (core waterbirds, taxid 3073812) at the time of initial analysis were downloaded using NCBI Datasets v17.1.0^70^ using the datasets download taxon commands with options –dehydrated –exclude-atypical –include genome –reference – mag exclude 3073812, and then downloaded with the datasets rehydrate command. This resulted in 89 seabirds, which were combined with our A9 and C90 reference genomes, for a total of 91 genomes to be aligned with ProgressiveCactus v2.9.7^71^. Seqkit v2.7.0^72^ was used to replace any ambiguous nucleotides with N. A phylogenetic tree was generated with Mashtree v1.4.6^73^, and the R package *ape*^74^ was used to convert it into a fully bifurcated tree.

### Variant calling

Variant calling was performed using GATK v. 4.2.1 as per our Nextflow script^75,76^. Four individuals were removed: PP6, PP19, YEP14, and N23-25, three due to low sequencing output, and one due to high heterozygosity indicative of sample mixing. We filtered variants using bcftools v1.18^77^ to remove genotypes missing in more than 20% of remaining samples, and a GQ of less than 20. For some downstream analyses, a minor allele count >= 2 was used. We imputed and phased samples using SHAPEIT v5.1.1^78^ for ancestral reconstruction graph construction.

### Ancient DNA variant calling

We downloaded the aDNA sample reads from Cole et al.^37^ from NCBI Sequence Read Archive; *M. a. richdalei* (SRR14902321) and *M. a. waitaha* (SRR14902318, SRR14902319). AdapterRemovalV2^79^ was used to identify and remove adapters and filter out poor-quality reads. Kraken2 v2.1.0^80^ was used to map reads to the Kraken PlusPF database to identify potential contaminants, common amongst aDNA studies^81^. Potential contaminant genomes were downloaded from NCBI (see Table S10), and concatenated with the putative ancestral basal penguin genome, Anc05, created by ProgressiveCactus above. We performed competitive mapping of the reads using bwa v0.7.19-r1273^82^ with the aln algorithm. We then performed three iterative mapping rounds to rescue reads, reducing mismatch penalties each time. All reads that did not map to the Anc05 genome were removed. Next, we used bamUtil v1.0.15^83^ to remove two bases from both ends of all reads to remove likely degraded basepairs^84^. ATLAS^85^, modified by for memory optimisation and resume functionality (https://github.com/jguhlin/ATLAS), was used to estimate errors and call SNPs. This comprehensive method was performed to reduce reference bias by mapping to a putative ancestral genome rather than our species of interest^86^ and rescue reads by iteratively reducing thresholds for unmapped reads^87,88^. Sequence locations were lifted over to the modern genome with halLiftOver from progressiveCactus^89,90^. Finally bcftools norm was used to normalise the resulting VCF files.

### Individual and per-population statistics

Per individual heterozygosity was calculated as (number of called sites - number of homozygous sites) /(number of called sites) as reported by the --het function in vcftools v1.15^91^. We utilised the --freq and --keep functions in vcftools to calculate allele frequencies per locus after subsampling 33 individuals each from Campbell Island and the Northern population to match the 33 individuals sampled from Enderby Island. Frequencies were folded to represent the minor allele frequency per locus, and monomorphic loci were removed. We repeated the subsampling 50 times for Campbell Island and the Northern population, to calculate mean and standard deviations per 0.05 frequency bin to generate a folded allele frequency spectrum per population. We utilised PLINK v1.09^92^ (https://www.cog-genomics.org/plink/) to infer runs of homozygosity (ROH) per individual, with key requirements that a window contain 30 SNPs, and excluding runs of less than 300kb (all parameter choices detailed in the Jupyter notebook). The sum of all identified runs of homozygosity was divided by the assembled genome size (1.36 Gb) to infer fraction of the genome in runs of homozygosity (FROH) as a metric of individual inbreeding.

### Phylogenetics

Phylogenetic analysis was conducted using BEAST v2.7.8^93^. Species trees were estimated from SNP alignments mapped to the reference genome for all individuals (excluding those removed due to sequencing failure), together with ancient DNA (aDNA) SNPs. In total, 400 genomic regions were analysed: 80 sets each derived from confidently aligned regions overlapping *M. a. waitaha* and *M. a. richdalei*, and a further 240 regions from extant yellow-eyed penguin (modern) populations. The latter were lifted over using halLiftOver v2.2^90^ from progressiveCactus v2.9.7^89^. Confident *M. a. waitaha* regions ranged from 400 to 1,801 bp, while *M. a. richdalei* regions spanned 400 to 5,101 bp. Regions from extant yellow-eyed penguins were uniformly 2 kbp in length and contained at least two, but fewer than 22, SNPs (2.5 × the standard deviation of binned SNP counts, rounded down).

Divergence dating was performed on 50 loci using the multispecies coalescent model implemented in StarBeast3 v1.2.1^42^, and using bModelTest v1.3.3^94^ to average over site and substitution models. These 50 loci each have over 20 parsimony informative sites in total, and at least one among the extant *Megadyptes antipodes*, and were randomly selected such that (i) at least tenloci were present in *waitaha*, (ii) at least tenloci were present in *richdaeli*, and (iii) the 30 loci contained more than three parsimony informative sites among the extant *antipodes*.

Divergence dates were estimated using five secondary calibrations among outgroup species from Cole et al.^37^ (as shown in Figure 3 and Table 1). All taxon ages were fixed at age 0, except for the aDNA samples from *M. a. waitaha* and *M. a. richdaeli*, which were respectively fixed at 750 and 3500 years before present^37^.

We assigned a fossilised birth-death prior to the species tree^45^, where the net diversification rate was sampled from a log-normal (μ=-2.5, σ=0.5) distribution, extinction fraction from Beta (α=3, β=1), and sampling proportion from Beta (α=1, β=5). Under this prior distribution, the expected tree height was 10.1 million years, with a 95% interquartile range of (0.29, 35.5). We used a strict and relaxed molecular clock with the rate drawn from a log-normal prior distribution, with a mean in real space of 0.007 substitutions per site per million years based on Cole et al.^37^, and a standard deviation of 1.25. The relaxed clock also had relative branch rates, each independently drawn from a log-normal prior distribution (mean of 1 in real space, standard deviation of 0.4 in log space). The mean effective population size (Ne) was estimated from a log-normal prior with a mean in real space of 0.01 million years and a broad standard deviation of 1.25. An Ne of 0.01 million years translates to an Ne of 1428 individuals, under the assumption of seven year generations. This estimate of Ne is much greater than the population size of *Megadyptes antipodes* (<3000 individuals) but much smaller than outgroups (>1 million rockhopper penguins (*Eudyptes*)). The Ne of each species is therefore drawn from an inverse gamma prior, with a shape of two, and a mean specified above. Additional phylogenies were built from SNPs, with no intervening reference sequence, using SNAPPER v1.1.5^39^ without time calibration (Figure 2b). Our BEAST 2 XML files, which contain the full model descriptions, are available on Github (https://github.com/GenomicsAotearoa/Hoiho).

### Population statistics

To test for signs of historical and recent migration, we ran BA3 v3.0.5.7^95^ on a random selection of 10,000 SNPs for eight million iterations with three million burn-in iterations for both all local regions and for the three main subspecies.

### Ancestral recombination graph construction

To construct the ancestral recombination graph, we first inferred the ancestral state of each position in the ProgressiveCactus ancestral penguin genome Anc05, which is basal to all penguins, by aligning against positions in the ancestral waterbird genome (Anc05) using a custom tool we developed, halAncestralAlleles (https://github.com/jguhlin/hal). We then inferred ARGs across the contigs at least 1Mb in size, and fewer than 25% unspecified ancestral alleles, in the A9 genome, spanning 1.13 Gbp and accounting for 83.24% of the genome length, using tsinfer v0.4.1^96^ with mismatch ratio of 1, and recombination rate of 2.5e-08, then dated with tsdate v0.2.3^96^ and using the tskit extend haplotypes functionality^97-100^.

### Genome-wide association studies

We fitted a generalised linear mixed-model profile likelihood ratio test (GLMM-PLRT) to perform a GWAS scan. This approach combines a logistic GLMM null^101^ fitted by Laplace-REML^102^ with an per-SNP projected score test transformed into a calibrated perturbation LRT and evaluated with a Gamma tail probability^103^. This is implemented with JAX v0.7.0^104^. We tested for both RDS and YPGV presence phenotypes within the Northern subspecies, and also ran the GWAS analyses for these phenotypes for all three subspecies. In each GWAS, we accounted for population structure by inferring genome-wide relatedness between individuals from their ancestral recombination graph with tskit^38^. We also fitted fixed effects of sex and individual inbreeding as inferred from FROH. Our

## Data availability

Genomic data generated in this project can be found in the Aotearoa Genomic Data Repository under doi: [pending]. All tools and scripts to generate the data are available at our github repository: https://github.com/GenomicsAotearoa/Hoiho.

## Author contributions

**Conceptualization**: JLG, JRW, AWS, JG, PSW, PKD. **Funding acquisition**: JLG, JRW, AWS, JG, PSW. **Sample Collection and Metadata**: JRW, MJY, TW, HS, JW, BA, TTR, LT, JVZ, MA, JW, HST, LSA. **PCR Tests**: JRW. **Necropsies**: JRW, HST, LSA, KM, SH. **Subspecies naming**: HL, PSW. **Population Genetic Analyses**: JG, AWS. **Phylogenetics**: JG, JD. **Writing–original draft**: JG, JRW, JD, JLG, AWS, CEG, PKD. **Writing–review, editing, final draft**: All Authors.

## Acknowledgements

We thank the Whenua Hou Komiti, and the four Papatipu Rūnaka (representing the Kaitiaki Roopū ki Murihiku), including Te Rūnaka o Waihopai, Te Rūnaka o Oraka-Aparima, Te Rūnaka o Hokonui, and Te Rūnaka o Awarua, for their input and kōrero during the planning, consultation and approval stages. We thank Kate McInnes, Bruce McKinlay, and Julia Reid from the New Zealand Department of Conservation. We thank the Dunedin Wildlife Hospital for providing resources for this project.

## Supplementary information

**Supplementary Figures:** Figures S1 - S6.

**Supplementary Tables:** Tables S1 - S10.

