## Supplementary Figures for "Population genomics of yellow-eyed penguins uncovers subspecies divergence and candidate genes linked to respiratory distress syndrome"

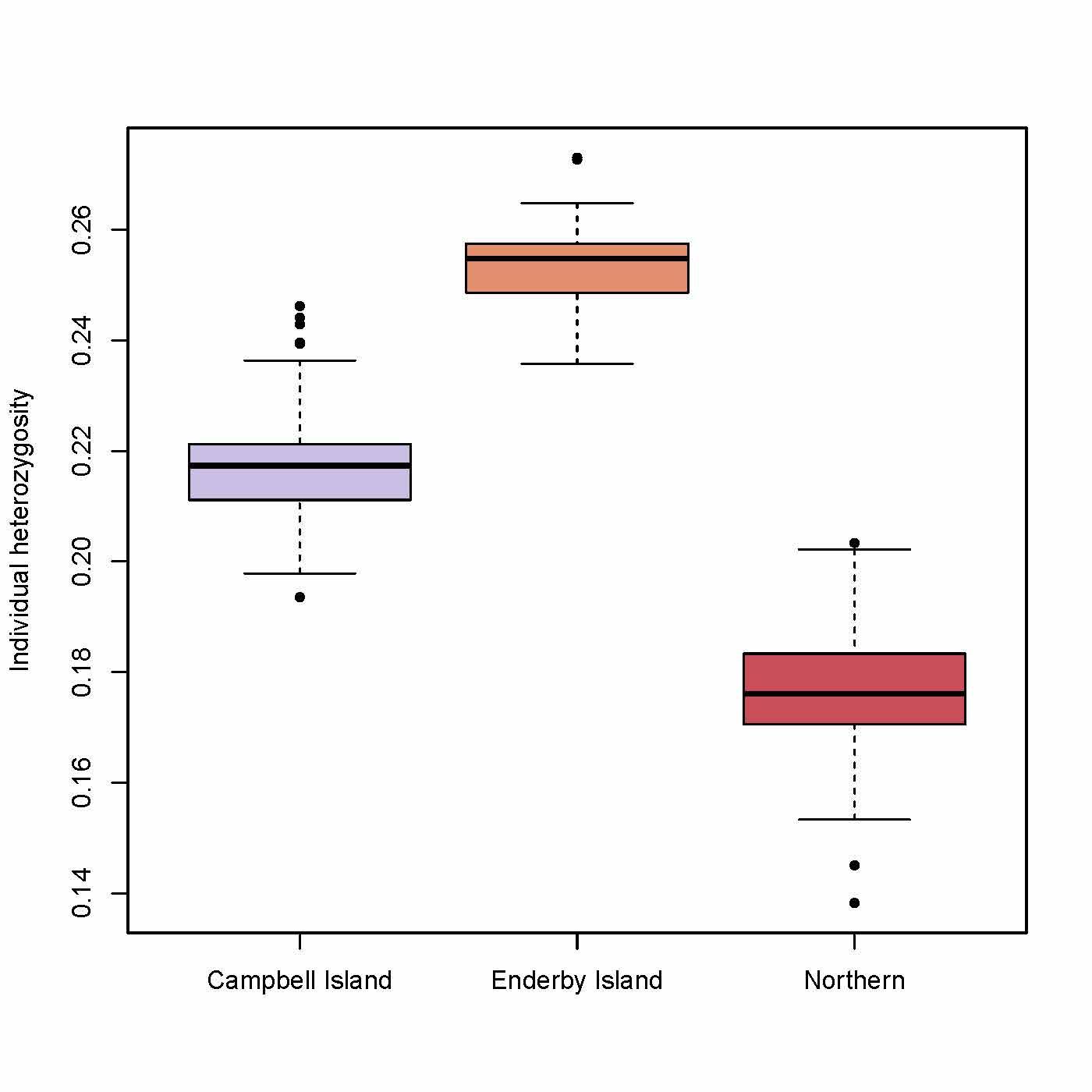

**Figure S1**. Individual heterozygosity at variable sites per subspecies, following quality filtering and excluding singletons per site. Campbell Island - 63 individuals, 1,232,200 SNPs; Enderby Island - 33 individuals, 1,058,683 SNPs; Northern subspecies - 153 individuals, 1,444,663 SNPs. Note that all individuals are mapped to the genome assembly of a Northern individual.

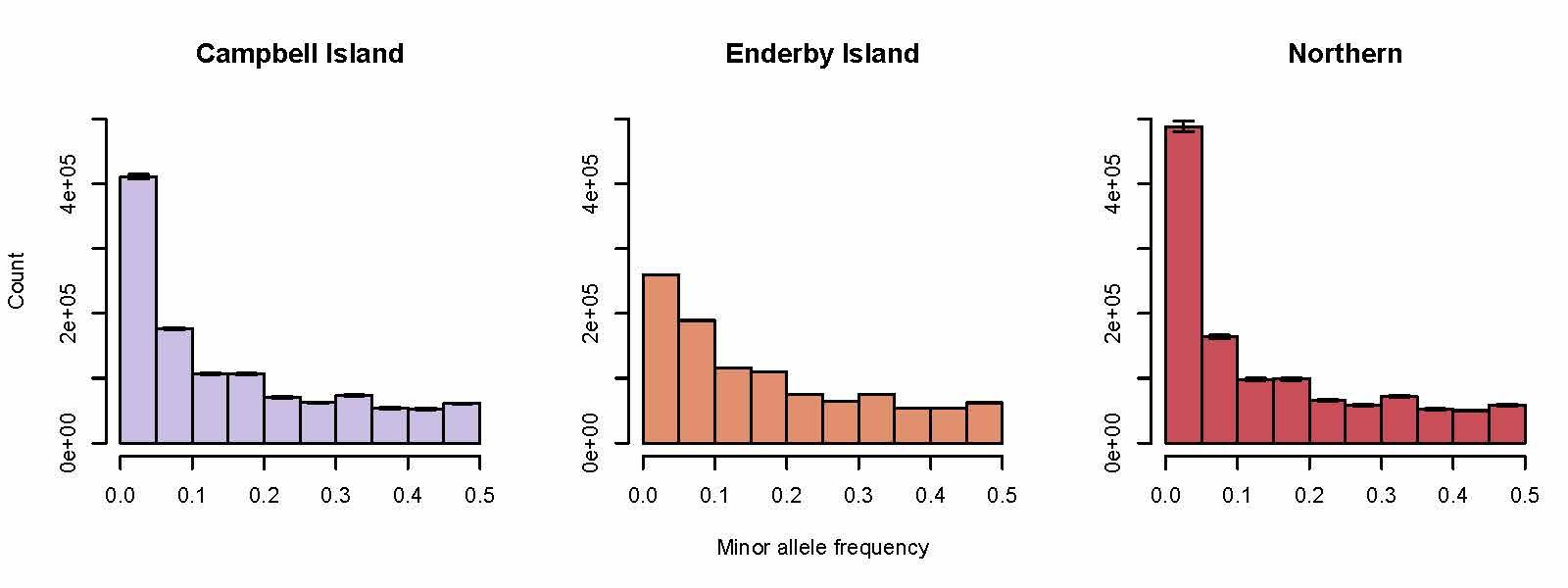

**Figure S2**. Folded allele frequency spectrum for variable sites per subspecies, following quality filtering and excluding singletons per site. Bin heights for Campbell Island and Northern represent mean values across 50 replicates when subsampling 33 individuals at random (to match the total number of individuals sampled for Enderby). Error bars are calculated per subspecies across these 50 replicates.

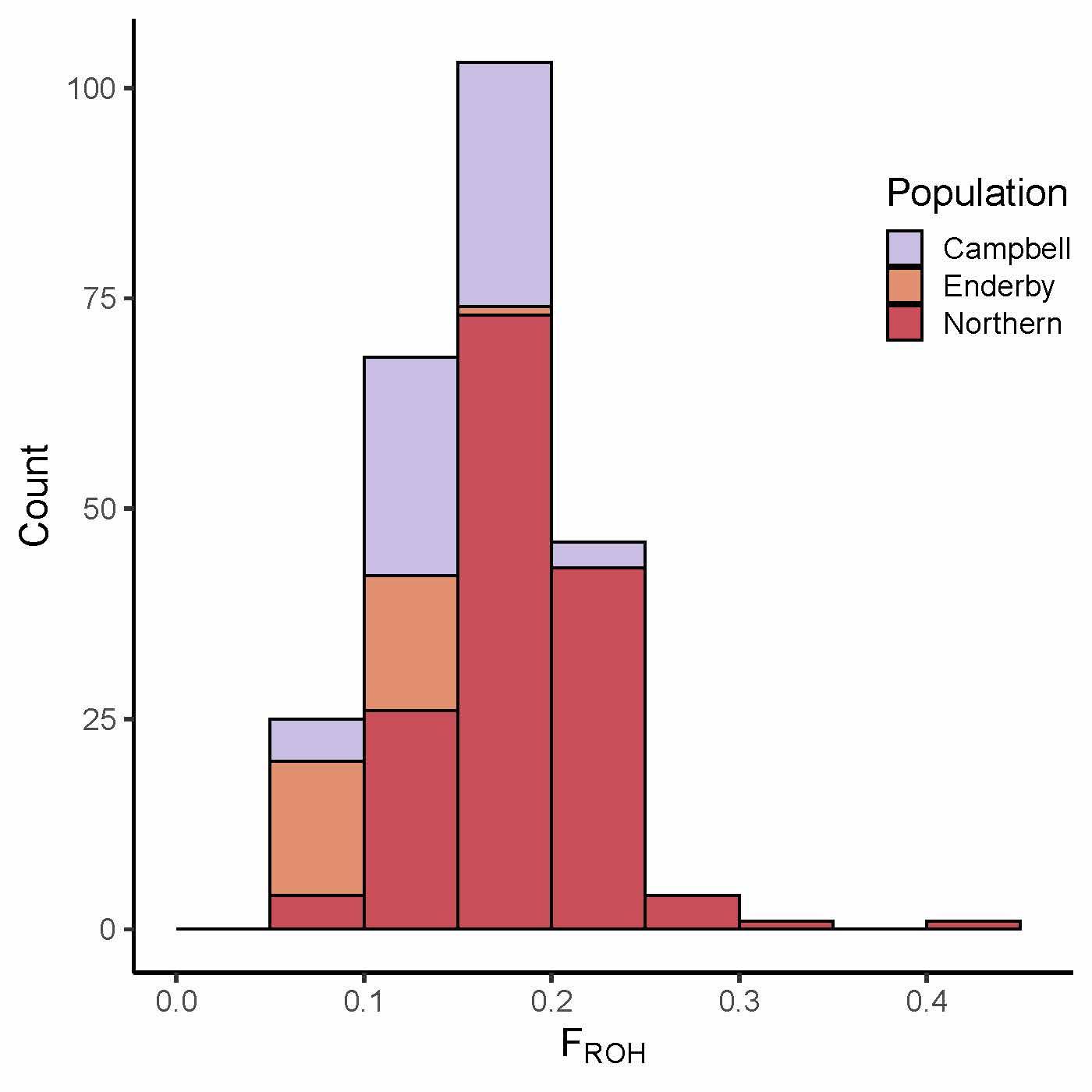

**Figure S3**. Distribution of inbreeding metrics for individuals sampled from Campbell Island, Enderby Island and Northern. Inbreeding calculated as F_ROH_, the sum of runs of homozygosity (ROH) of >300kb divided by the genome assembly size (1.36 Gb).

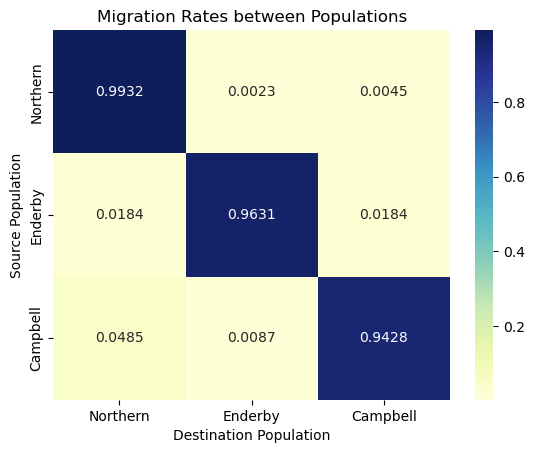

**Figure S4**. Migration between the three major populations as determined by BA3-SNP. Std deviations, per row then per column, are reported as follows: 0.0038, 0.0022, 0.0031, 0.0126, 0.175, 0.0182, 0.0132, 0.0061, 0.0142).

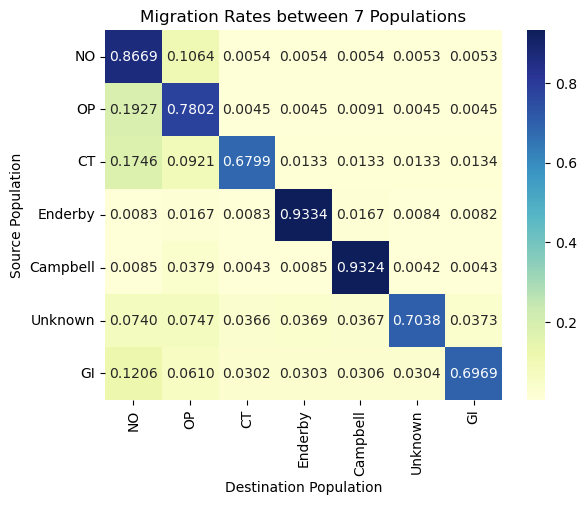

**Figure S5**. Migration rates between the 7 populations, this includes Northern population sub-locations. For brevity, standard deviations are not included.

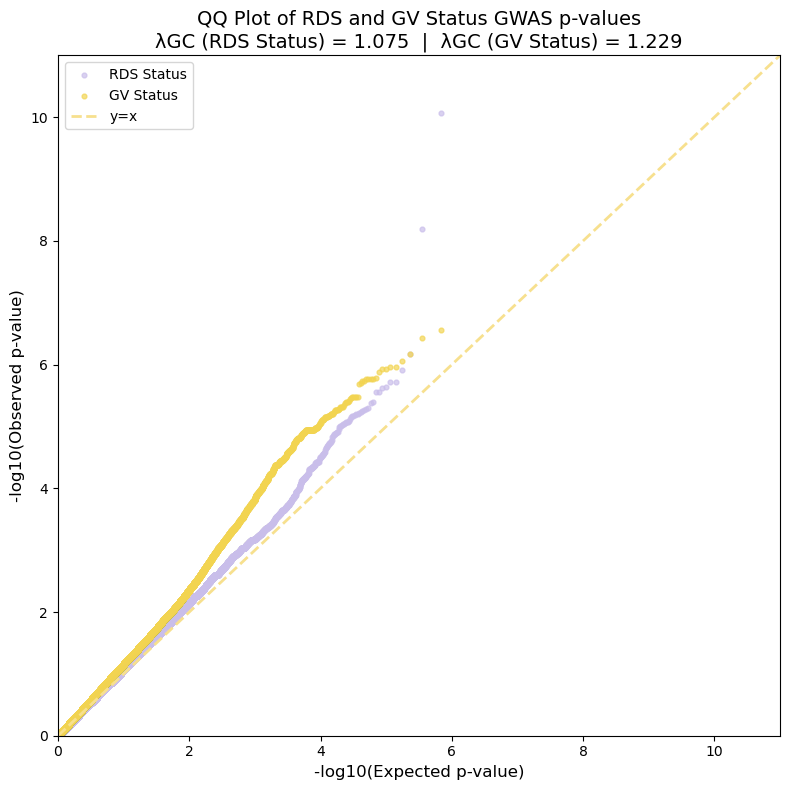

**Figure S6**. QQ-Plot for our GWAS analysis for RDS Status and GV status. Lambda GC for both analyses is listed at the top. Lambda GC is calculated as the ratio of the median observed chi-squared test statistic to the expected median under the null hypothesis.λ1
